# Structural basis of main proteases of coronavirus bound to drug candidate PF-07321332

**DOI:** 10.1101/2021.11.05.467529

**Authors:** Jian Li, Cheng Lin, Xuelan Zhou, Fanglin Zhong, Pei Zeng, Yang Yang, Yuting Zhang, Bo Yu, Xiaona Fan, Peter J. McCormick, Rui Fu, Yang Fu, Haihai Jiang, Jin Zhang

## Abstract

The high mutation rate of COVID-19 and the prevalence of multiple variants strongly support the need for pharmacological options to complement vaccine strategies. One region that appears highly conserved among different genus of coronaviruses is the substrate binding site of the main protease (M^pro^ or 3CL^pro^), making it an attractive target for the development of broad-spectrum drugs for multiple coronaviruses. PF-07321332 developed by Pfizer is the first orally administered inhibitor targeting the main protease of SARS-CoV-2, which also has shown potency against other coronaviruses. Here we report three crystal structures of main protease of SARS-CoV-2, SARS-CoV and MERS-CoV bound to the inhibitor PF-07321332. The structures reveal a ligand-binding site that is conserved among SARS-CoV-2, SARS-CoV and MERS-CoV, providing insights into the mechanism of inhibition of viral replication. The long and narrow cavity in the cleft between domains I and II of main protease harbors multiple inhibitor binding sites, where PF-07321332 occupies subsites S1, S2 and S4 and appears more restricted compared with other inhibitors. A detailed analysis of these structures illuminated key structural determinants essential for inhibition and elucidated the binding mode of action of main proteases from different coronaviruses. Given the importance of main protease for the treatment of SARS-CoV-2 infection, insights derived from this study should accelerate the design of safer and more effective antivirals.

## Introduction

Since its discovery in Dec 2019, cases of novel severe acute respiratory syndrome coronavirus 2 (SARS-CoV-2)-infected pneumonia have rapidly continued to emerge, with the current case count close to 250 million and a case mortality rate ∼3.4%, causing huge economic and social loss to the world.^[1, 2]^ There have been several coronaviruses in human history that are pathogenic to humans, among which two were associated with severe respiratory disease outbreaks: SARS-CoV (severe acute respiratory syndrome coronavirus first emerged in Guangdong China in 2002) and MERS-CoV (Middle East respiratory syndrome coronavirus first detected in Saudi Arabia in 2012).^[3-5]^ Genomic sequencing data showed that SARS-CoV-2 shares 79.6% sequence identity with SARS-CoV.^[1, 6-8]^ As SARS-CoV and SARS-CoV-2 successively emerged and COVID-19 continues spreading throughout the globe, developing broad spectrum drugs remains an urgent and unmet clinical need in the treatment and prevention of COVID-19 infections.

Viral enzymes and proteins of CoVs that are involved in coronavirus replication are potential drug targets for COVID-19. In particular, the main protease (M^pro^ or 3CL^pro^), which cleaves the replicase polyproteins at 11 sites, is one of the most attractive targets for numerous classes of small molecule inhibitors for the development of drugs against coronavirus infections.^[9, 10]^ M^pro^ is highly conserved among coronaviruses, and the substrate-binding site in M^pro^ also share several common features.^[6]^ Because M^pro^ has no human homolog, M^pro^ inhibitors should be highly specific to SARS-CoV-2 and have minimal side effects.^[11]^ Unfortunately, although several peptidomimetic covalent inhibitors of M^pro^ have been reported, few candidates has progressed into clinical trials.^[6, 12-14]^

We previously screened an in-house small molecule library and identified shikonin as a noncovalent inhibitor for SARS-CoV-2 and SARS-CoV M^pro^ in vitro, with a half-maximum inhibitory concentration (IC50) of 1.57 μM and 7.89 μM, respectively.^[15, 16]^ Besides, crystal structures of SARS-CoV-2 M^pro^ with antineoplastic drug carmofur and natural products baicailin and baicalein have also been solved recently^[17, 18]^. Despite the progress made in understanding the origin of human coronaviruses as well as the prediction and prevention of the emerging pandemics, we still lack effective and safe drugs and therapies to combat the global pandemic caused by coronavirus.^[5, 19, 20]^

In March 2021, Pfizer has initiated a Phase 1 clinical trial of a novel antiviral therapeutic agent against SARS-CoV-2. The clinical candidate, PF-07321332, is the first orally administered coronavirus-specific protease inhibitor, which has shown potency in vitro anti-viral activity against SARS-CoV-2, as well as activity against other coronaviruses.^[21]^ As a protease inhibitor, PF-07321332 binds to the viral enzyme and can block the activity of the protease that the coronavirus needs to reproduce itself. This kind of inhibitors have effectively treated other viral pathogens, such as HIV and hepatitis C virus. As there is currently no orally administered therapy approved for the post-exposure or pre-emptive treatment of COVID-19, PF-07321332 could be an encouraging therapeutic with potential for use in the treatment of COVID-19, as well as potential use to address future coronavirus threats.

In this study, we aim to explore the molecular basis for the small molecule inhibitor PF-07321332 targeting M^pro^ of coronaviruses. We found that PF-07321332 potently inhibits the enzymatic activity of SARS-CoV-2 M^pro^. We then determined the crystal structures of complexes of main protease of SARS-CoV-2, SARS-CoV and MERS-CoV bound to the inhibitor PF-07321332, revealing a novel binding mode of M^pro^. Structural comparison with reported M^pro^-inhibitor complex structures provides an insight into the mechanism of M^pro^ inhibition by a small molecule inhibitor and a framework for small molecule drug discovery.

## Results

### Inhibitory activity of PF-07321332 against SARS-CoV-2 M^pro^

We purified the M^pro^ of SARS-CoV-2 as previously reported^[15]^. A Fluorescence Resonance Energy Transfer (FRET) assay was employed to determine the inhibitory activity of PF-07321332 against SARS-CoV-2 M^pro^. Ebselen was used as a control with an IC50 value of 0.365 μM, which is similar to previous studies^[22] [23]^. The results showed that PF-07321332 has a potent inhibition with the IC_50_ value against the M^pro^ of SARS-CoV-2 being 0.023 μM (Fig. 1), which is much lower that recently reported covalent inhibitors such as boceprevir, leupeptin, and carmofur ^[24-26]^.

**Fig. 1.**
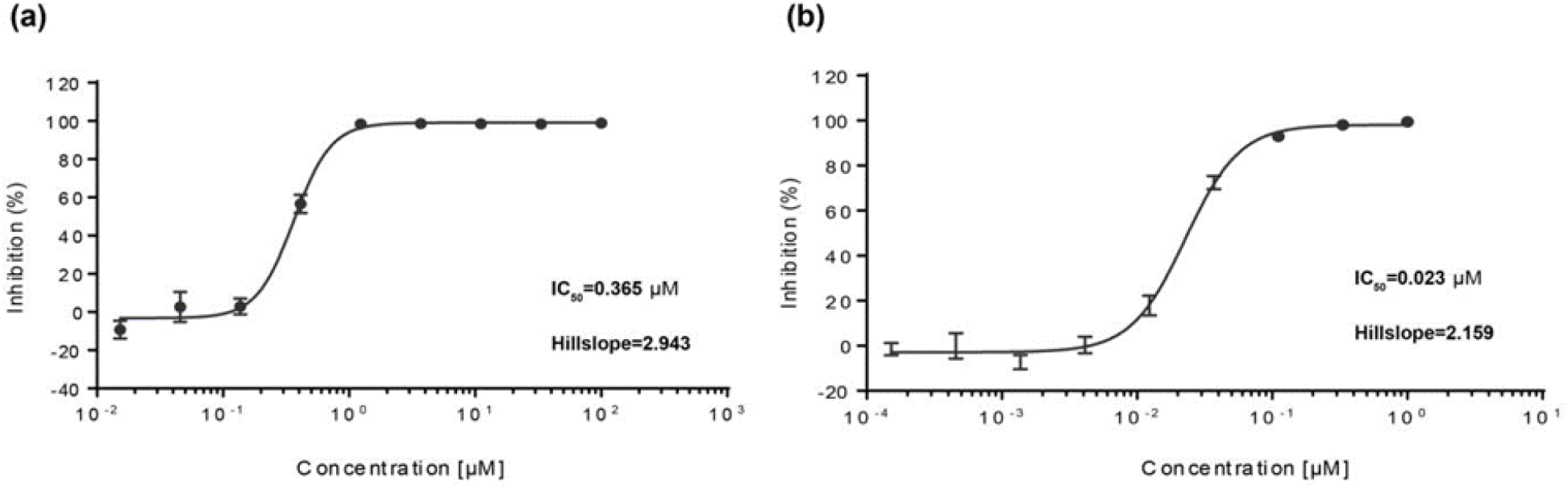
Enzymatic inhibition of SARS-CoV-2 Mpro. (a) Inhibition of ebselen against SARS-CoV-2 Mpro; (b) Inhibition of PF-07321332 against SARS-CoV-2 Mpro. SAR-CoV-2 Mpro was preincubated in the reaction buffer with various concentrations of PF-07321332 at room temperature for 30 min before reacting with the FRET substrate. Ebselen was used as a control. The IC50 was calculated using the GraphPad Prism software.

### Inhibitory mechanisms of PF-07321332 against SARS-CoV-2 M^pro^

In order to figure out the inhibitory mechanisms of PF-07321332, we determined the crystal structure of SARS-CoV-2 M^pro^ in complex with PF-07321332 (PFM^pro^-Co) at 1.5 Å resolution by co-crystallization method (Table 1). By comparison with the apo structure of SARS-CoV-2 M^pro^ at pH 7.5, the root mean square deviation (RMSD) of equivalent C□ positions between apo Mpro and PFM^pro^-Co is ∼ 0.985 Å (Fig. S1). As shown in Fig. 2, the M^pro^ molecule in the complex structure forms a homodimer and contains three domains, namely domain I (residues 10-99), domain II (residues 100-184), and domain III (residues 201-303). PF-07321332 can be found in both protomer A and protomer B (Fig. 2a). Specifically, PF-07321332 binds to the active site situated in the cleft between domains I and II of M^pro^ in an extended conformation and occupies subsites S1, S2 and S4 of SARS-CoV-2 M^pro^ (Fig. 2b). The electron density map unambiguously shows that nitrile carbon of PF-07321332 forms a C-S covalent bond with the sulfur atom of catalytic residue C145 (Fig. 2c). The imine nitrogen of the thioimidate moiety occupies the oxyanion hole and forms hydrogen bonds with the backbone NH of C145 and the oxygen from a water molecule for stabilization. Besides the typical covalent interaction, PF-07321332 forms multiple non-covalent interactions with the active site. According to the Berger and Schechter nomenclature, PF-07321332 consists of five moieties, namely P1-P4 and P1’. As shown in Fig. 2d and 2e, PF-07321332 contains a γ-lactam ring at P1 position just before the warhead nitrile group. The lactam ring inserts into the S1 subsite with the oxygen and nitrogen atoms of the lactam ring forming hydrogen bonds with the N_ε_2 of H163 and carboxy group of E166, respectively. Further, a hydrogen bond is also found between the amide nitrogen at the P1 moiety and the main-chain carbonyl oxygen of H164. The P2 position contains a dimethylcyclopropylproline (DMCP) moiety which inserts into the S2 subsite and mainly forms hydrophobic interactions, similar with the previously reported compound boceprevir ^[25]^. PF-07321332 presents a tert-leucine residue at P3 position. However, limited interactions are observed between P3 moiety of PF-07321332 and S3 subsite of M^pro^. PF-07321332 displays a trifluoromethyl group at P4 position. The amide nitrogen at P4 position forms a hydrogen bond with the main chain carbonyl oxygen of E166, while the trifluoromethyl group forms additional hydrogen bonds by interacting with the nitrogen atom of Q192 and a water molecule. Thus, PF-07321332 occupies the active site of SARS-CoV M^pro^ by covalently binding to C145 and non-covalently interacting with conserved residues including Cy145, H163, H164, E166 and Q192. The crystal structure of M^pro^-PF-07321332 complex (PFM^pro^-So) has also been solved at 1.9 Å resolution by soaking (Table 1). By superimposition of PFM^pro^-Co and PFM^pro^-So, we found that the binding mode of PF-07321332 with SARS-CoV-2 Mpro was highly similar with the RMSD being 0.781 Å over the 523 best-aligned Cα atoms (Fig. S2).

**Table 1.**
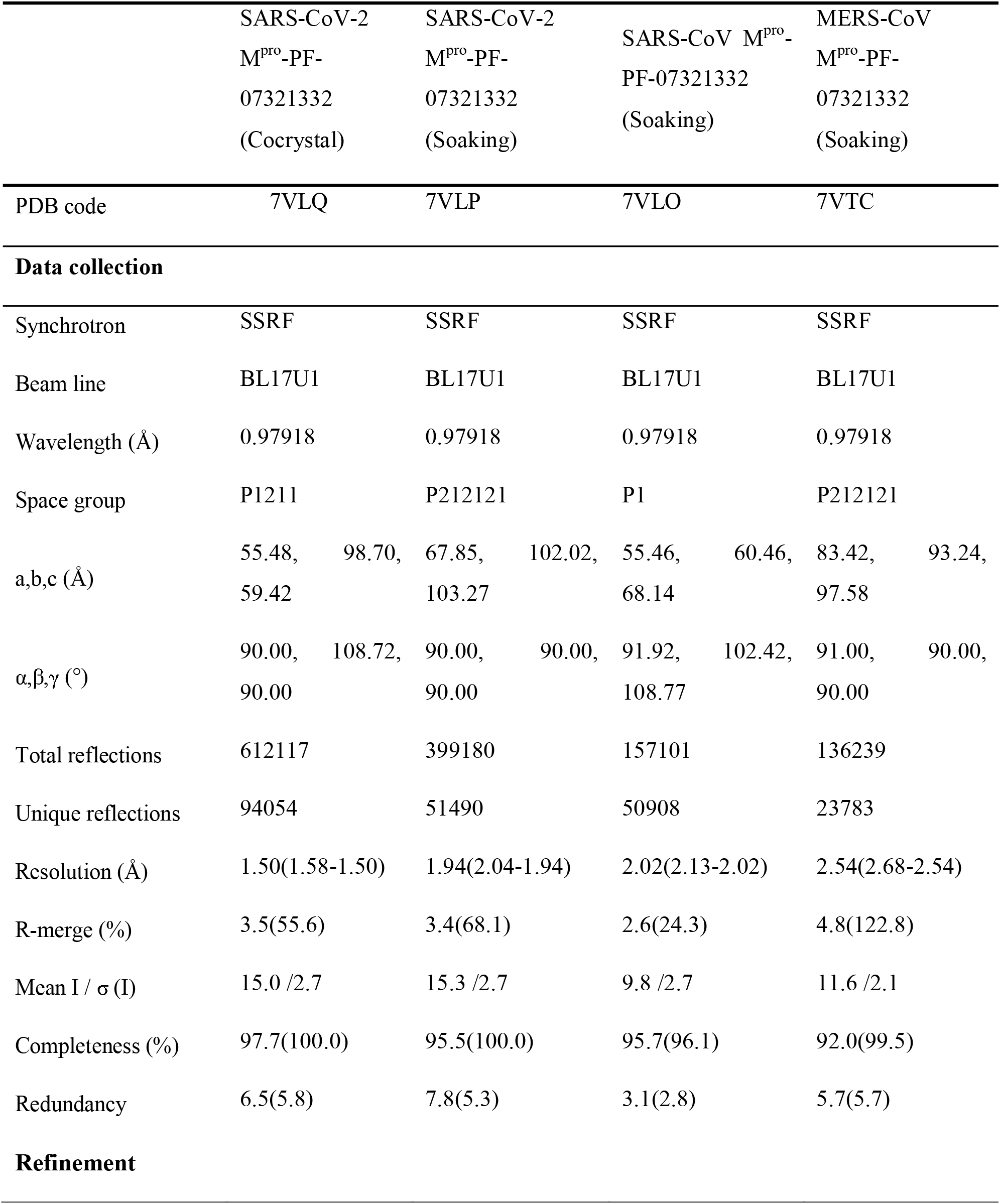
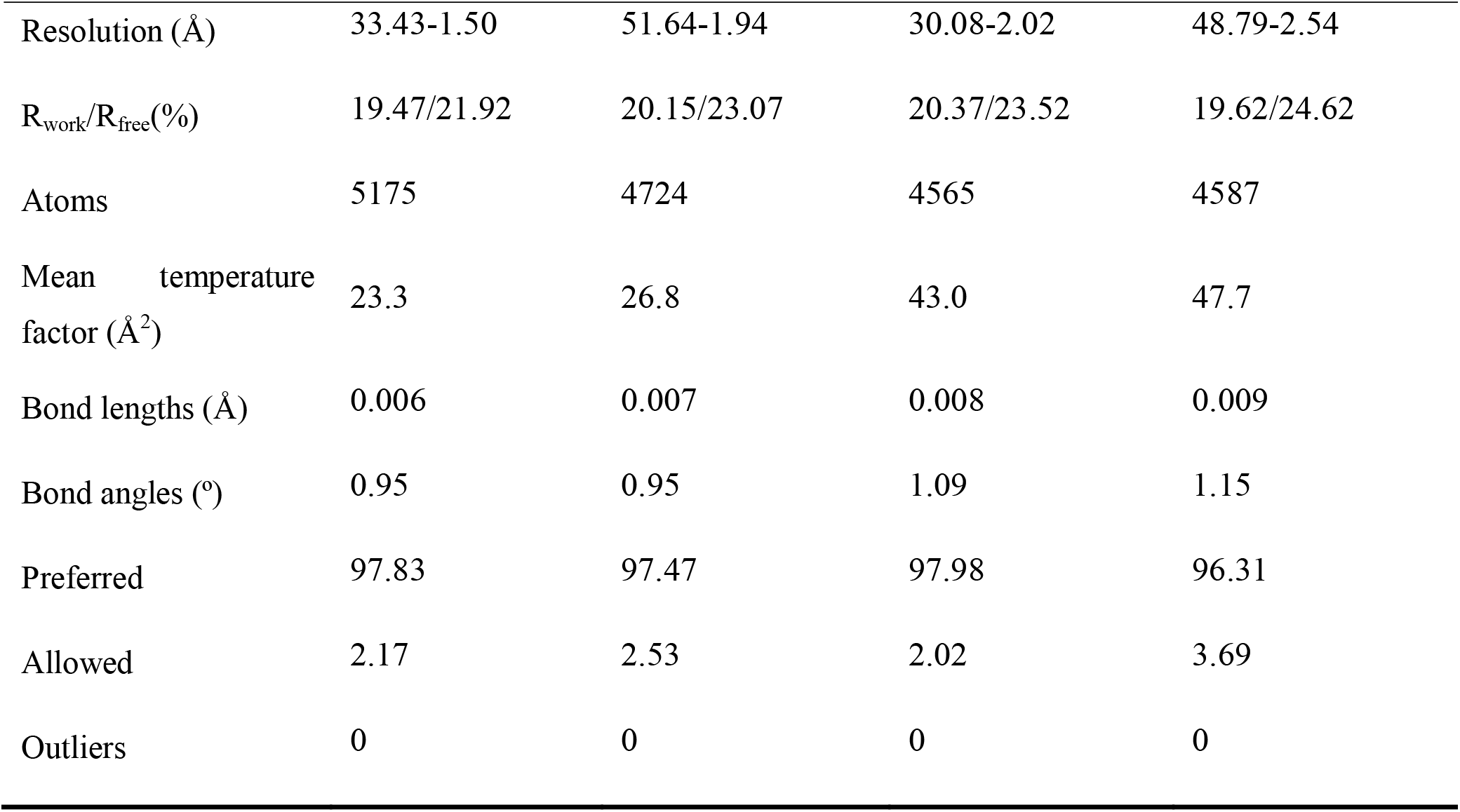
Statistics for data processing and model refinement of M^pro^-PF-07321332

**Fig. 2.**
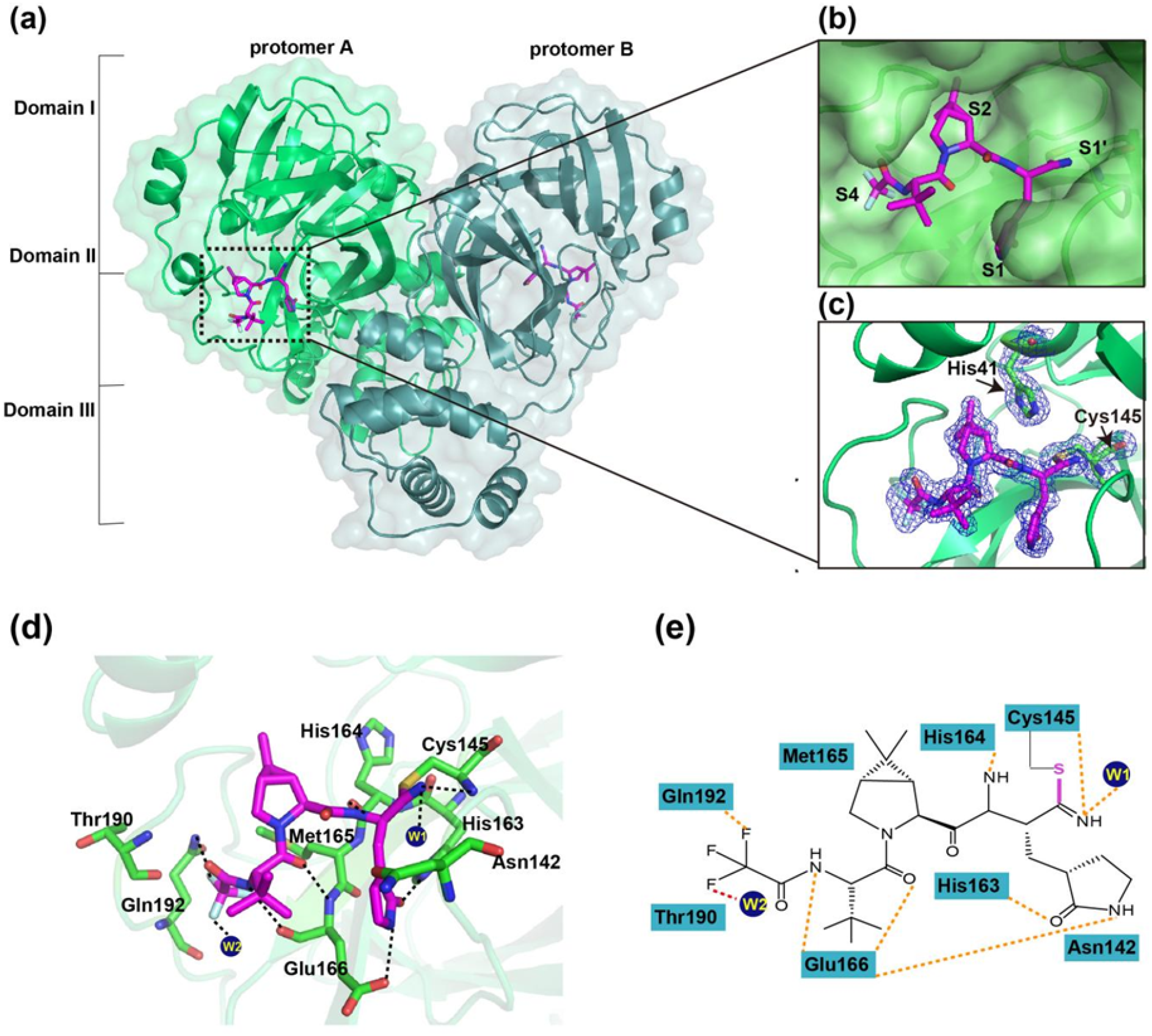
Crystal structure of SARS-CoV-2 Mpro in complex with PF-07321332. (a) Overall structure of SARS-CoV-2 Mpro in complex with PF-07321332. The three domains and two protomers of Mpro are labeled. The substrate-binding pocket is located within the black dotted box. PF-07321332 is shown as sticks with the carbon atoms in magentas, oxygen atoms in bright red, and nitrogen atoms in blue, and flfluorine atom in palecyan. (b) An enlarged view of the substrate-binding pocket. PF-07321332 forms a covalent bond with C145. The substrate-binding subsites (S1’, S1, S2 and S4) are labeled. (c) A C-S covalent bond forms between the S_γ_atom of C145 and the nitrile carbon of PF-07321332. The 2Fo_™_Fc density map contoured at 1.0_σ_i shown as blue mesh. (e) The detailed interaction in the complex structure is shown with the residues involved in inhibitor binding (within 3.5 Å) displayed as sticks. W1 and W2 represent the water molecules. Hydrogen bonds interactions are shown as black dashed lines. (e) Schematic interaction between PF-07321332 and Mpro. Hydrogen bonds interactions are shown as orange dashed lines.

### Crystal structures of PF-07321332 in complex with SARS-CoV and MERS-CoV M^pro^

We also determined the crystal structure of PF-07321332 in complex with SARS-CoV and MERS-CoV M^pro^ at 2.0 Å and 2.5 Å resolution (Table 1), respectively. PF-07321332 displays a highly similar conformation in the substrate-binding site of SARS-CoV, MERS-CoV, and SARS-CoV-2 M^pro^ even though the orientation of each moiety of PF-07321332 has slight differences (Fig. 3a and 3b). Like the case with SARS-CoV-2 M^pro^, PF-07321332 fits into the S1, S2 and S4 subsites, and forms a covalent bond with the catalytic residue cysteine as expected (Fig. 3c and 3d). In addition, several amino acid residues, including F140, C145, H163, H164, E166 and Q192, in the protease form hydrogen bond interactions with the inhibitor in the SARS-CoV M^pro^-PF-07321332 complex. The key residues of MERS-CoV M^pro^ interacting with PF-07321332 is highly conserved (Fig. 3e-3h). These observations are consistent with the fact that the structure of M^pro^ in the coronaviruses is highly conserved and that PF-07321332 may be a potent inhibitor with broad-spectrum potential to defeat diseases caused by various coronaviruses.

**Fig. 3.**
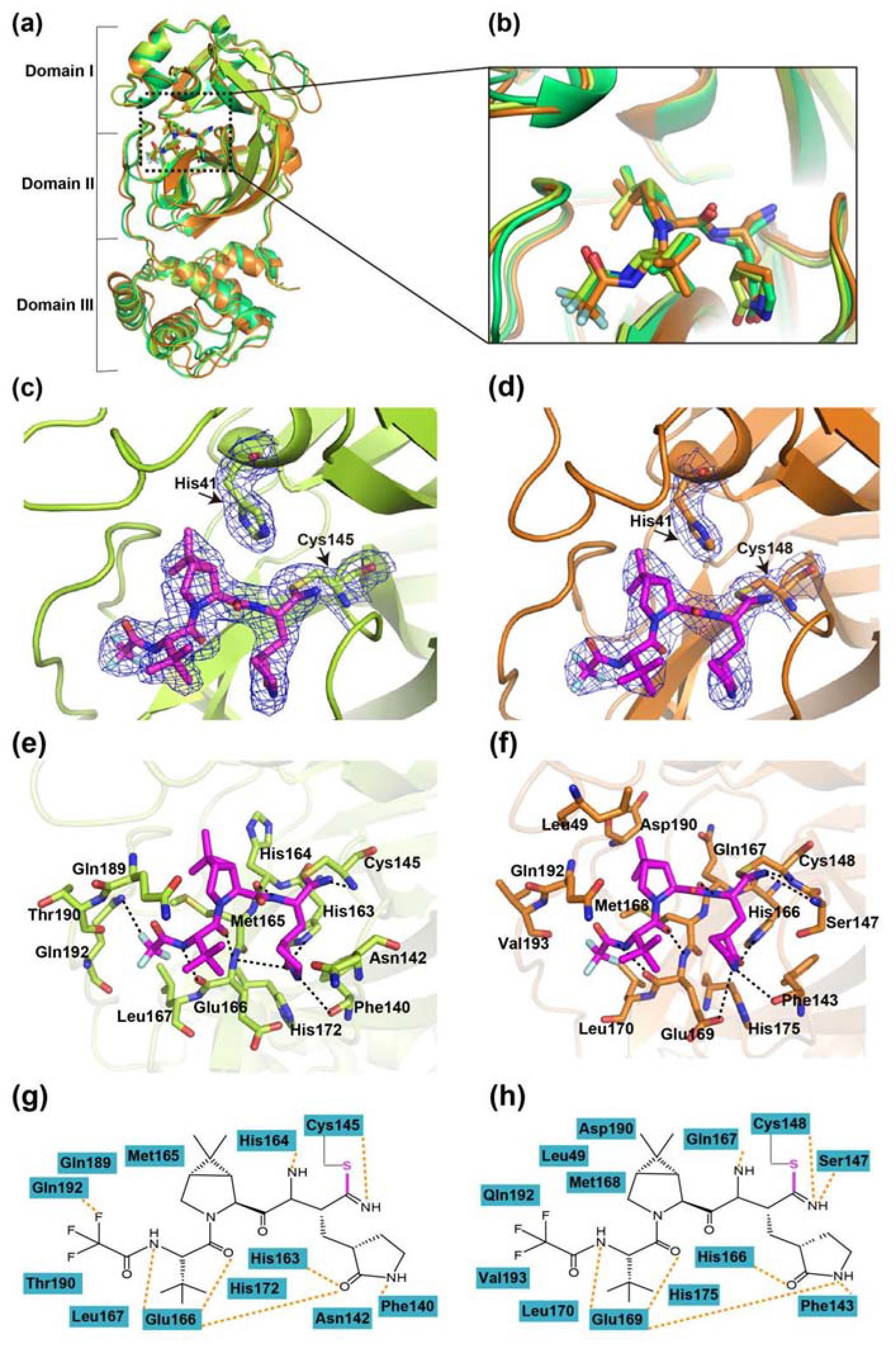
Crystal structures of SARS-CoV and MERS-CoV Mpros in complex with PF-07321332. (a) Structural alignment of CoV Mpros complexed with PF-07321332 with SARS-CoV-2 Mpro-inhibitor complex in limegreen, SARS-CoV Mpro-inhibitor complex in limon, and MERS-CoV Mpro-inhibitor complex in orange. (b) An enlarged view of the substrate-binding pocket. (c and d) A C-S covalent bond forms between C145 of SARS-CoV Mpro (c) or MERS-CoV Mpro (d) and the nitrile group of PF-07321332. The 2Fo-Fc density map contoured at 1.0_σ_is shown as blue mesh. (e and f) The detailed interaction in the complex structure is shown with the residues of SARS-CoV Mpro (e) or MERS-CoV Mpro (f) involved in inhibitor binding (within 3.5 Å) displayed as sticks. Hydrogen bonds interactions are shown as black dashed lines. (g and h) Schematic interaction between PF-07321332 and SARS-CoV Mpro (g) or MERS-CoV Mpro (h). Hydrogen bonds interactions are shown as orange dashed lines.

### Structural comparison of PF-07321332 with other covalent inhibitors in complex with SARS-CoV-2 M^pro^

We also compared the structures of the recently identified covalent inhibitors complexed with SARS-CoV-2 M^pro^. All these small molecules form a C-S covalent bond with the catalytic residue cysteine (Fig. 4) and display a lactam ring at P1 position which fits well with the S1 subsite except carmofur and boceprevir. The amide nitrogen of the lactam ring can form hydrogen-bond interactions with the ketonic oxygen of Glu166 or Phe140, while the oxygen atom form hydrogen-bond interactions with backbone NH of Glu166 or Nε2 of His163. Subsite S2 of SARS-CoV-2 M^pro^ appears to prefer hydrophobic interactions and all these inhibitors display a hydrophobic group at P2 position, such as isopropyl and the dimethylcyclopropylproline (DMCP) group. However, the P3 position of these inhibitors does not fit well with the S3 pocket. Indeed, structural optimization may be conducted in the future to investigate more suitable groups for this subsite. The binding pattern of PF-07321332 is similar with that of boceprevir at P4 position but differs at substitution of F atoms and formation of hydrogen bonds with Q192. Therefore, structures of inhibitors in complex with M^pro^ reported in this study and other studies, will provide the structural basis for the development and optimization of more potent drugs against SARS-CoV-2 infection.

**Fig. 4.**
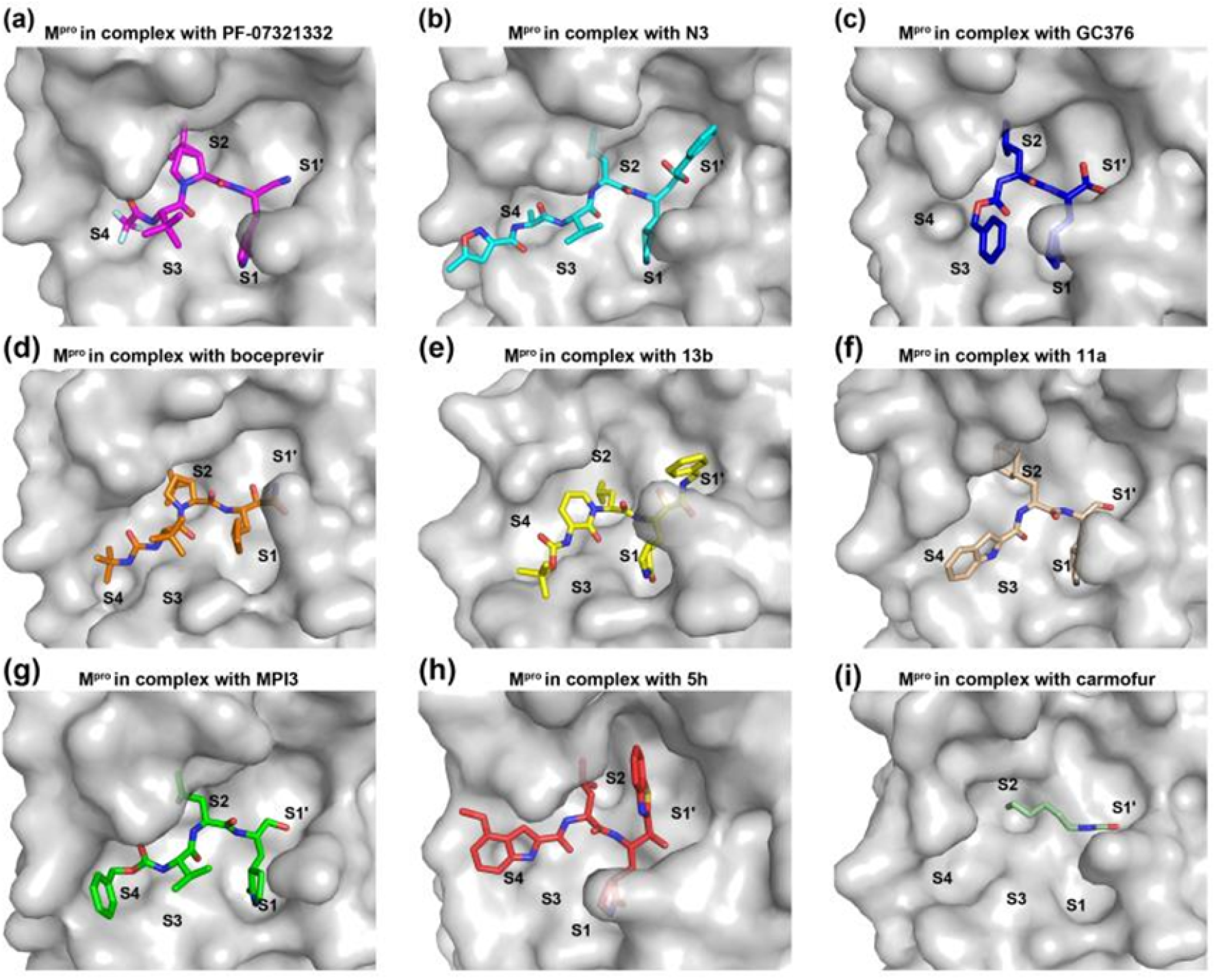
Comparison of the binding modes of different inhibitors targeting SARS-CoV Mpro. The binding pockets of PF-07321332 (a, PDB: 7VLQ), N3 (b, PDB: 6LU7), GC376 (c, PDB: 7D1M), boceprevir (d, PDB: 7BRP), 13b (e, PDB: 6Y2F), 11a (f, PDB: 6LZE), MPI3 (g, PDB: 7JQO), 5h (h, PDB: 7JKV) and carmofur (i, PDB: 7BUY) bound to SARS-CoV-2 Mpro are shown, respectively. Mpros are show as surface in gray and the inhibitors are shown as sticks.

## Discussion

There is an urgent need to develop effective drugs as the novel coronavirus pandemic continues to wreak havoc on human society. M^pro^ is a promising drug target for its vital role in viral replication, high conservation among all coronavirus, and has no homolog in humans. Although a great number of inhibitors have been found to show inhibitory activity at M^pro^, most of these are not potent enough and not highly bioavailable following oral administration, which functionally limits its clinical application to hospitalized patients with relatively advanced disease. Thus, potent orally bioavailable antiviral drugs for treatment of SARS-CoV-2 infection are urgently needed. PF-07321332 is a SARS-CoV-2 M^pro^ inhibitor developed by Pfizer in clinical trial ^[27]^. In this study, we found PF-07321332 is a potent inhibitor with a IC_50_ being 0.023 μM. Though this value is a little higher than MPI3, the most potent inhibitor of SARS-CoV-2 M^pro^ so far ^[28]^, but lower than most reported inhibitors including 11a (IC_50_=0.053□μM) ^[29]^, 13b (IC_50_=0.67□μM)^[30]^, boceprevir (IC_50_=8.0□μM) ^[24]^ and GC376 (IC_50_=0.15□μM) ^[24]^, which implicates its huge potential to treat COVID-19 clinically. In addition, the main advantage of PF-07321332 with respect to other inhibitors of SARS-CoV-2 M^pro^ is the possibility of oral administration, a feature that could dramatically facilitate the treatment of COVID-19.

We also solved the crystal structure of PF-07321332 in complex with M^pro^ of three deadly coronoviruses (SARS-CoV-2, SARS-CoV and MERS), which is of great importance for structure-based drug development. The complex structures indicate that PF-07321332 covalently bound to the catalytic cysteine of M^pro^, and formed multiple hydrogen bonds with conserved residues within the active site. By comparing with other covalent inhibitors, PF-07321332 employs a unique binding pattern to SARS-CoV-2 M^pro^. Further structure-based optimization of such covalent inhibitors will help generate drugs against current COVID-19 pandemic with high efficiency and broad-spectrum.

Besides PF-07321332, another orally bioavailable drug candidate, masitinib, has been demonstrated to show inhibitory effect against SAR-CoV-2 M^pro^. Up to now, masitinib has been approved for treatment of mast cell tumors in dogs and evaluated in phase 2 and 3 clinical trials in humans for the treatment of cancer, asthma, Alzheimer’s disease, multiple sclerosis, and amyotrophic lateral sclerosis ^[31]^. However, unlike PF-07321332 as a covalent inhibitor, masitinib is a non-covalent inhibitor, which could complement the design of covalent inhibitors against SARS-CoV-2 main protease. Both covalent and non-covalent M^pro^ inhibitor will contribute to increase public health preparedness for potential future pandemic.

## Materials and Methods

### Expression and purification of human CoVs

The codon-optimized cDNAs for M^pro^ of SARS-CoV-2, SARS and MERS were synthesized fused with 6 His at the N terminus. Synthesized genes were subcloned into the pET-28a vector. The expression and purification of each main protease performed by a standard method described previously by our lab.^[15]^

### Enzymatic Assays

Fluorogenic substrates as a donor and quencher pair were synthesized. The IC_50_ values of the screening compounds against SARS-CoV-2, SARS and MERS main proteases were measured with a common protocol as the following: First, 1 μL of six human CoVs proteases (200 nM) was incubated with various concentrations of testing inhibitors at room temperature for 30 min in its reaction buffer (50 mM Tris 7.3,150mM NaCl, 1 mM EDTA) in a 384-well plate, and then FRET substrate was added to the reaction system. The reaction was monitored for 20 min, and the data was calculated at 10 min intervals by linear regression. The IC_50_ of compounds was determined by plotting the initial velocity against various concentrations of testing inhibitor by using the dose-response curve in GraphPad Prism software.

### Crystallization

Co-crystallization of SARS-CoV-2 with PF07321332(0.1 M HEPES pH 7.5, 20% w/v PEG 10000), SARS with PF07321332(0.1M HEPES pH7.5, 10%PEG 8000, 8% Ethylene glycol) and MERS with PF07321332(10% PEG 200, 0.1M Bis-Tris-propane pH9.0, 18% PEG 8000) were carried out at 20 °C using the hanging drop vapor-diffusion method, PF07321332 was added to according to a 3:1 molar ratio and the mixture was incubated for 30 min on ice, after 3-5 days, the complex crystals of human CoVs with PF07321332 were obtained.SARS-CoV-2-apo crystals were conducted using sitting-drop vapor diffusion method at 20°C. PF07321332 was soaked with crystals of SARS-CoV-2-apo(0.1 M HEPES sodium pH 7.5, 10% v/v 2-Propanol, 20% w/v PEG4000) within 24 h.

### Data collection, structure determination and refinement

The crystals were tailored with cryo-loop and then flash-frozen in liquid nitrogen to collect better X-ray data. All data sets were collected at 100 K on macromolecular crystallography beamline17U1 (BL17U1) at Shanghai Synchrotron Radiation Facility (SSRF, Shanghai, China). All collected data were handled by the HKL 2000 software package. The structures were determined by molecular replacement with PHENIX software. The program Coot was used to rebuild the initial model. The complete wanted data collection and statistics of refinement are shown in Table 1. Coordinates and structure factors for SARS-PF07321332, SARS-CoV-2-PF07321332 (soaking and Co-crystallization) and MERS-07321332 complexes have been deposited in the Protein Data Bank (PDB) under accession numbers 7VLO, 7VLP, 7VLQ and 7VTC, respectively.

## Acknowledgements

We would like to thank the Cryo-EM center of Southern University of Science and Technology for our Cryo-EM work and their help of Cryo-EM data collection. J.L. was supported by the Open Project of Key Laboratory of Prevention and treatment of cardiovascular and cerebrovascular diseases, Ministry of Education (No. XN201904), Gannan Medical University (QD201910), Jiangxi key research and development program (20203BBG73063) and Jiangxi “Double Thousand Plan”. J.Z. was supported by the Thousand Young Talents Program of China, the National Natural Science Foundation of China (grant no. 31770795; grant no. 81974514), and the Jiangxi Province Natural Science Foundation (grant no. 20181ACB20014). P.J.M was supported by the Foreign Talent project of Jiangxi Province. F.Y. was supported by Shenzhen Science and Technology Program (JCYJ20210324115611032 and KQTD20200909113758004). This work was also supported by Ganzhou COVID-19 Emergency Research Project (2020.17), Major science and technology programs of Ganzhou City (2020.67) and Ganzhou Zhanggong District COVID-19 prevention and control key research projects (2020.67).

## Author Contributions

J.L. and J.Z. initiated and supervised the project. C.L., X.Z., F.Z. and P.Z. crystallized the protein complexes, performed soaking experiments. J.L., C.L., X.Z., F.Z., P.Z. and J.Z. collected X-ray data, and solved and refined structures. B.Y., Y.Z. and H.J. performed docking and identified compounds to be tested in the initial screens and assisted enzymatic assay by FRET assay. H.J., P.J.M., Y.Y., X.F., Y.F. and R.F. assisted with design of experiments, project management, and interpretation of results.

