## Supplemental materials for "Structural basis of main proteases of coronavirus bound to drug candidate PF-07321332"

Figure S1

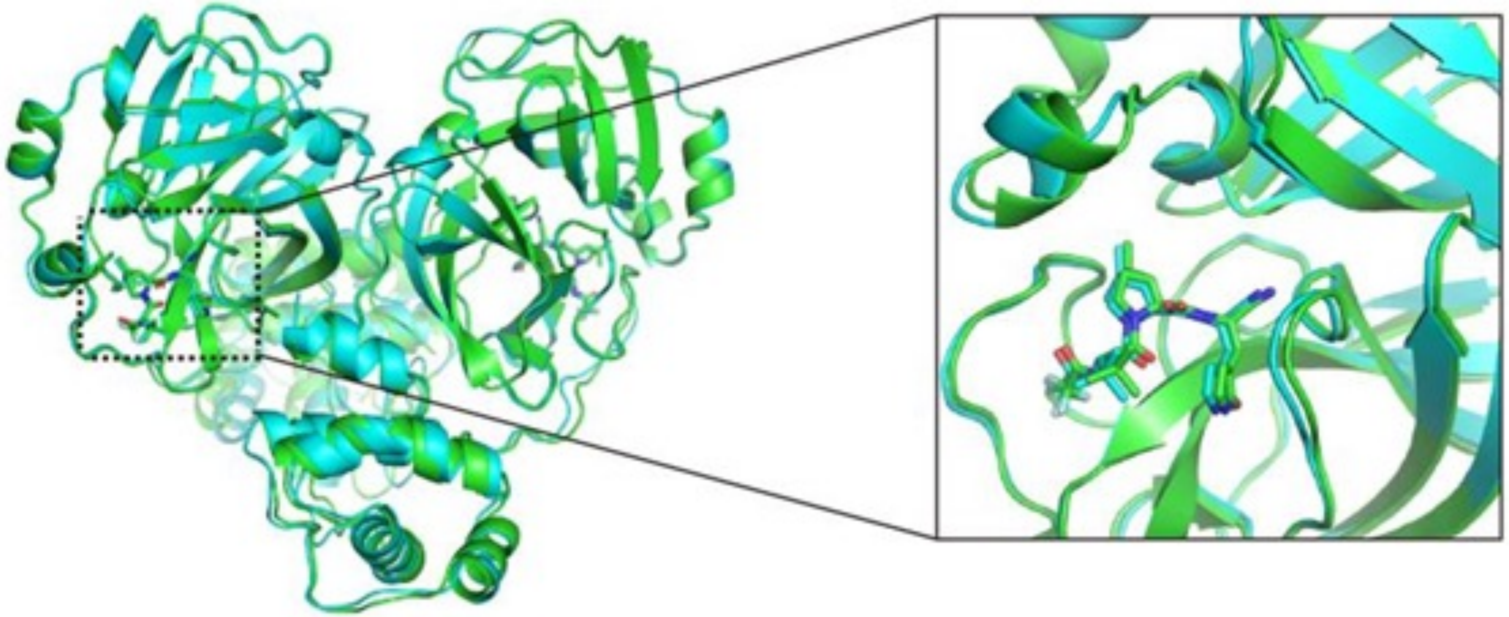

Fig. S1 Structural comparison of PF-07321332/SARS-CoV-2 $M^{pro}$ complexes obtained by cocrystallization (green) with soaking (cyan).

Figure S2

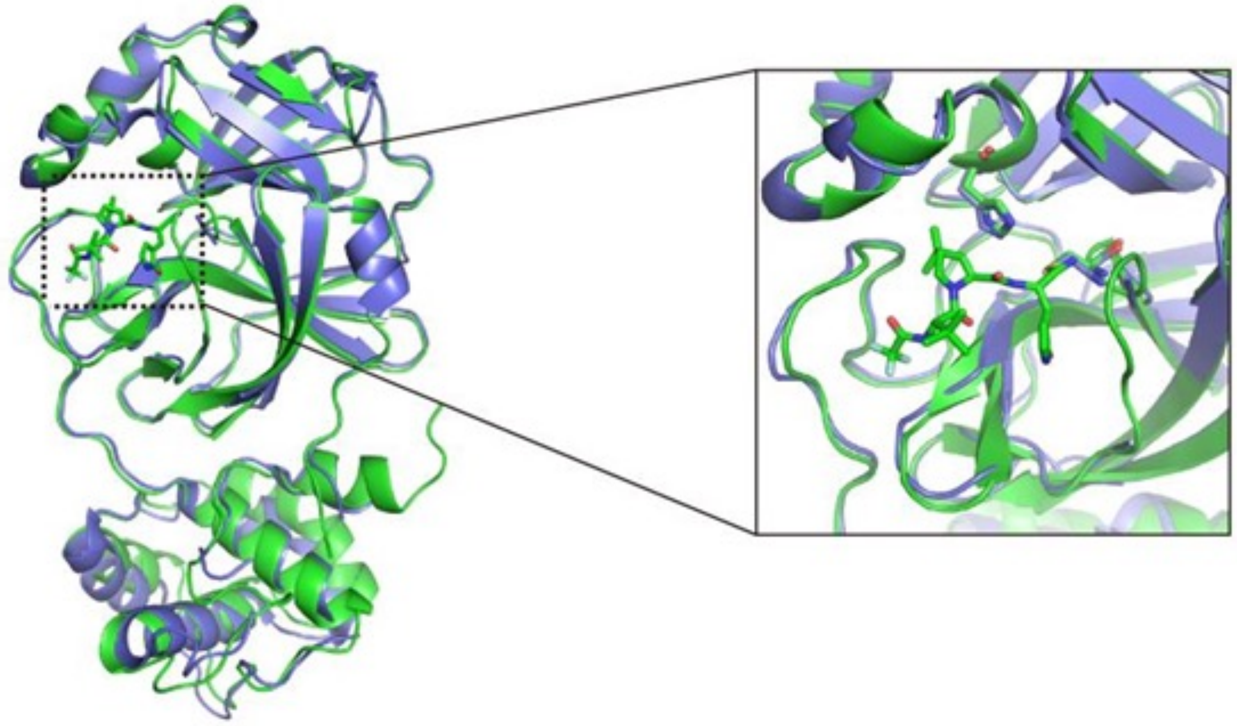

Fig. S2 Structural comparison of PF-07321332/SARS-CoV-2 $M^{pro}$ complexes obtained by cocrystallization (green) with apo $M^{pro}$ (slate).

Figure S3

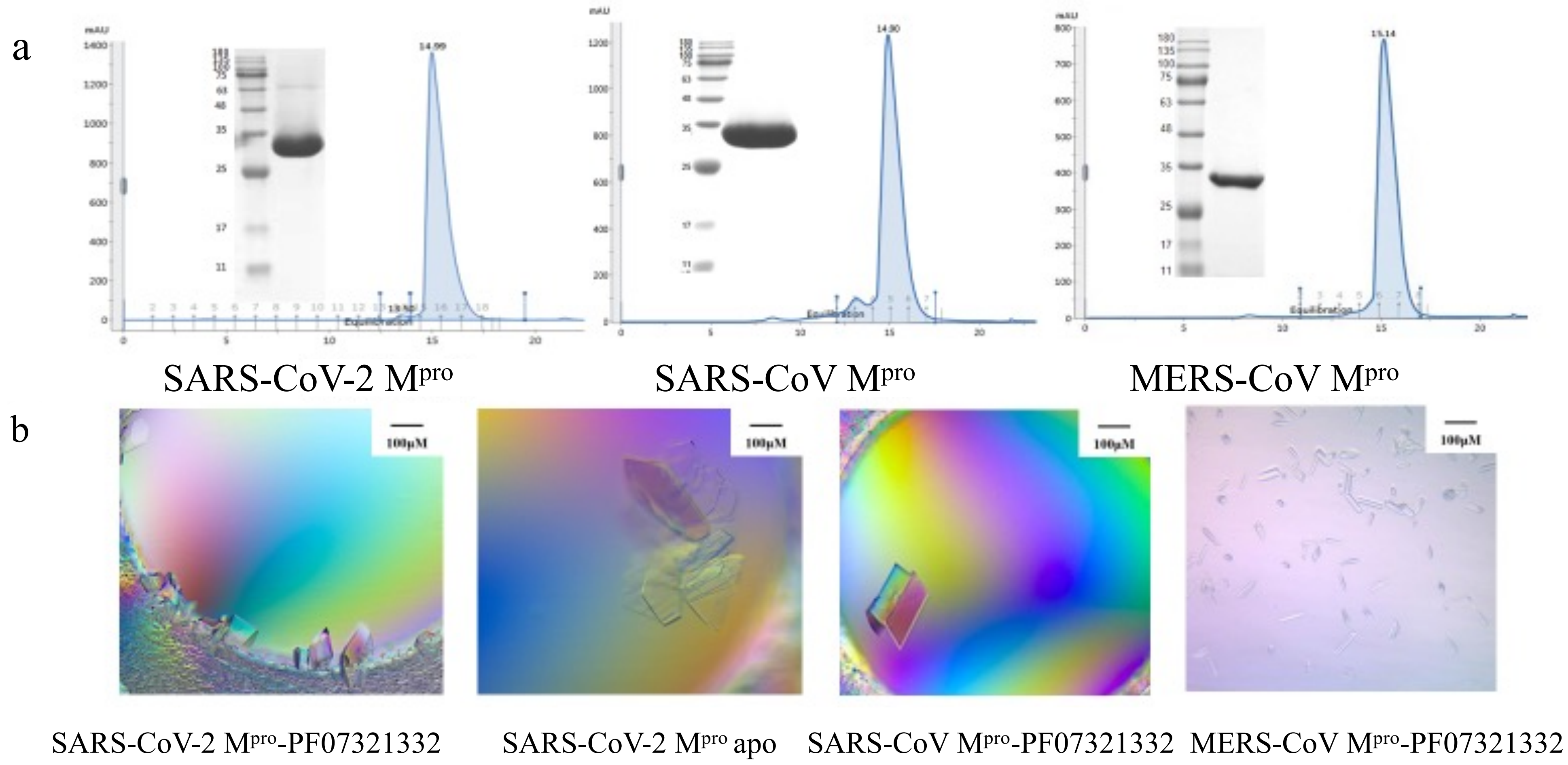


Fig. S3 a Purification of human CoVs . b Crystals of SARS-Cov-2 and human CoVs with PF07321332.
